## Supporting Information File 1 for "Elevated levels of lamin A promote HR and NHEJ-mediated repair mechanisms in etoposide-treated ovarian cancer cells"

**Table 1.**

**siRNA Sequence:**

| siRNA | Sequence Sequence |
| --- | --- |
| siRNA 1 | 5’GGUGGUGACGAUCUGGGCU3’ |
| siRNA 2 | 5’AACUGGACUUCCAGAAGAACAUC3’ |

**Table 2.**

**Antibodies:**

| **Name** | **Company** | **Dilution** |
| --- | --- | --- |
| Lamin A | Sigma-Aldrich | IF: 1:100 ; WB : 1:500 |
| Lamin B | SantaCruz | IF: 1:50 ; WB : 1:100 |
| γH2AX | EMD-Millipore | IF: 1:50 ; WB : 1:1200 |
| Rad51 | Abcam | IF: 1:50 ; WB : 1:500 |
| BRCA1 | EMD-Millipore | IF: 1:50 |
| BRCA2 | Thermo Fischer Scientific | IF: 1:50 |
| Ku70 | SantaCruz | IF: 1:50 ; WB : 1:1000 |
| PCNA | SantaCruz | IF: 1:50 ; WB : 1:500 |
| βActin | Sigma-Aldrich | WB : 1:1000 |
| Anti-BrdU | SantaCruz | IF : 1:50 |

**Table 3.**

**Primer Sequences:**

| **Primer** | **Sequence** |
| --- | --- |
| GAPDH | Forward: 5’- GAAGGTGAAGGTCGGAGTCAAC -3’  Reverse: 5’- CAGAGTTAAAAGCAGCCCTGGT -3’ |
| LA | Forward: 5’ -CGGTTCCCACCAAAGTTCA -3’  Reverse: 5’-CTCATCCTCGTCGTCCTCAA -3’ |
| LB | Forward: 5’- AAAAGACAACTCTCGTCGCAT- 3’  Reverse: 5’ -CCGCTTTCCTCTAGTTGTACG -3’ |
| Rad51 | Forward: 5'-TCTCTGGCAGTGATGTCCTGGA-3'  Reverse: 5'-TAAAGGGCGGTGGCACTGTCTA-3' |
| BRCA1 | Forward: 5'-CTGAAGACTGCTCAGGGCTATC-3'  Reverse: 5'-AGGGTAGCTGTTAGAAGGCTGG-3' |
| Ku70 | Forward: 5'-TGCCACAGGAAGAAGAGTTG-3'  Reverse: 5′-CTCTGGAGTTGCCATGATTT-3' |
| PIF1 | Forward: 5’- GGTAAGGTACACAGATTTGAGGC-3’  Reverse: 5’-CCCGAGACACCGATAAGTTTT-3’ |
| RIF1 | Forward: 5’-TGTTGGAGACTTTGGAAGACC-3’  Reverse: 5’-ACTTTGTACAGCCGAGGAAG-3’ |
| BRCA2 | Forward: 5’-TTCATGGAGCAGAACTGGTG-3’  Reverse: 5’-AGGAAAAGGTCTAGGGTCAGG-3’ |
| FGF2 | Forward: 5’-ACCCTCACATCAAGCTACAAC-3’  Reverse: 5’-AAAAGAAACACTCATCCGTAACAC-3’ |
| TLR2 | Forward: 5’-TGGTAGTTGTGGGTTGAAGC-3’  Reverse: 5’- GACAGAGAAGCCTGATTGGAG-3’ |
| BIRC3 | Forward: 5’-AATGCTTTTGCTGTGATGGTG-3’  Reverse: 5’-GCTTGAACTTGACGGATGAAC-3’ |
| THBS1 | Forward: 5’-CTCCCCTATGCTATCACAACG-3’  Reverse: 5’-AGGAACTGTGGCATTGGAG-3’ |
| PLK1 | Forward: 5’-ACAGTTTCGAGGTGGATGTG-3’  Reverse: 5’-GGTTGATGTGCTTGGGAATAC-3’ |
| XRCC2 | Forward: 5’-CAGTTGGTGAATGGCGTTG-3’  Reverse: 5’-CTACCTTCAAGTCGGGCAAG-3’ |
| POLQ | Forward: 5’-GCCAGGGTTCTCTATGCTTC-3’  Reverse: 5’-TCTTCAACTGCTTCCTCTTCC-3’ |
| MCM10 | Forward: 5’-AACCAGCCATCAAGTCCATC-3’  Reverse: 5’-TGGGCTCTCAACTTCACTTG-3’ |
| BRIP1 | Forward: 5’-GCTTAGCCTTACTTTGTTCTGC-3’  Reverse: 5’-TTTCACTTACGCCCTCATCTG-3’ |

**Supplementary Figures:**

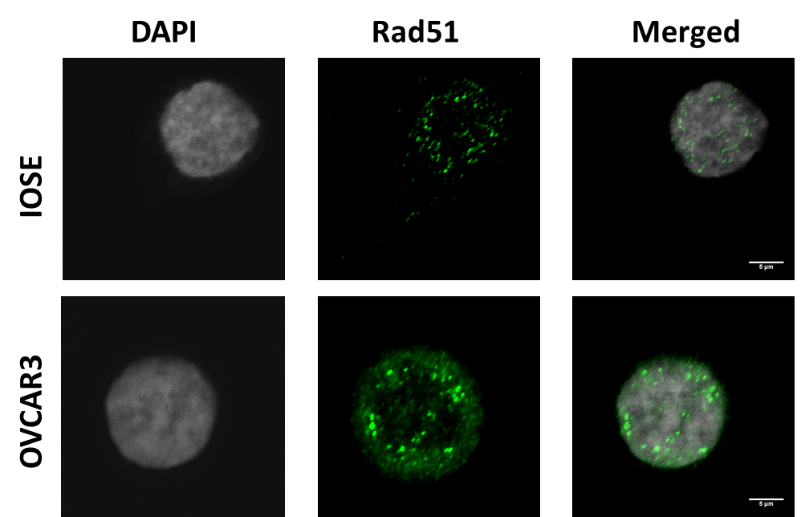

**Figure 1.** Confocal images of OVCAR3 and IOSE nuclei stained with Rad51. Magnification: 60X. Scale Bar: 5µm

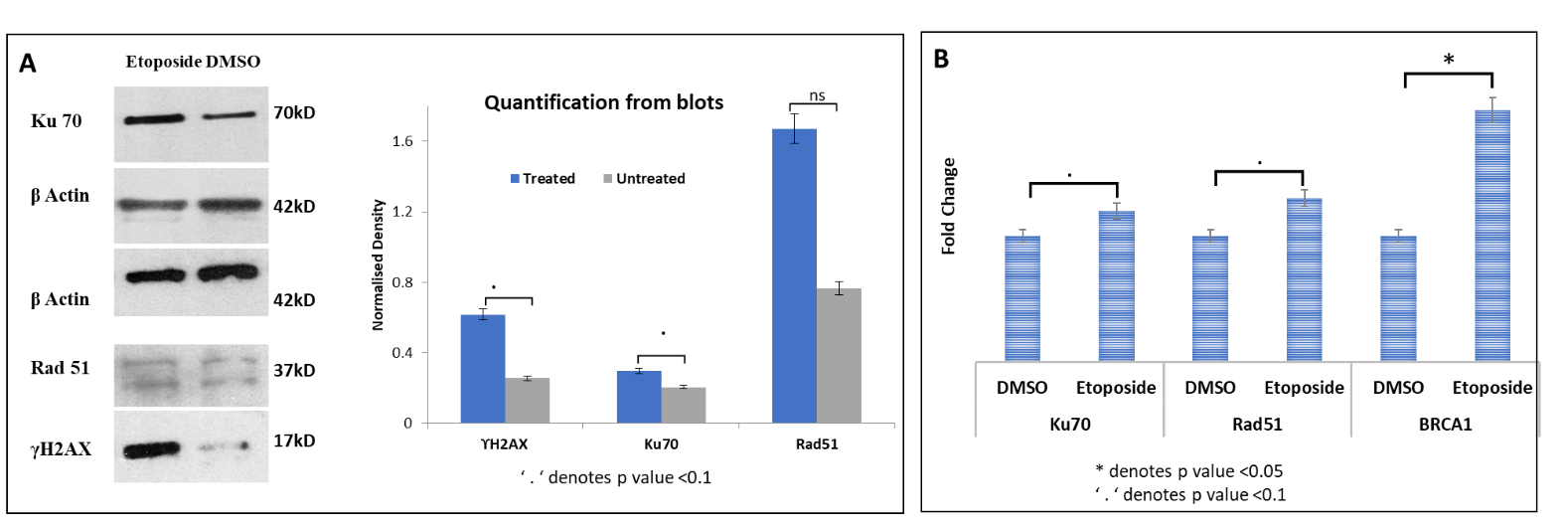

**Figure 2:** Western Blots showing the level of Ku70, Rad51 and γH2AX in etoposide treated and untreated OVCAR3 cells. β Actin is used as loading control. **D.** qPCR showing fold changes of DNA Damage repair proteins in OVCAR3 cells treated with etoposide. Error bar indicates standard error. * indicates p value to be <0.05, ‘ . ‘ indicates p value to be <0.1

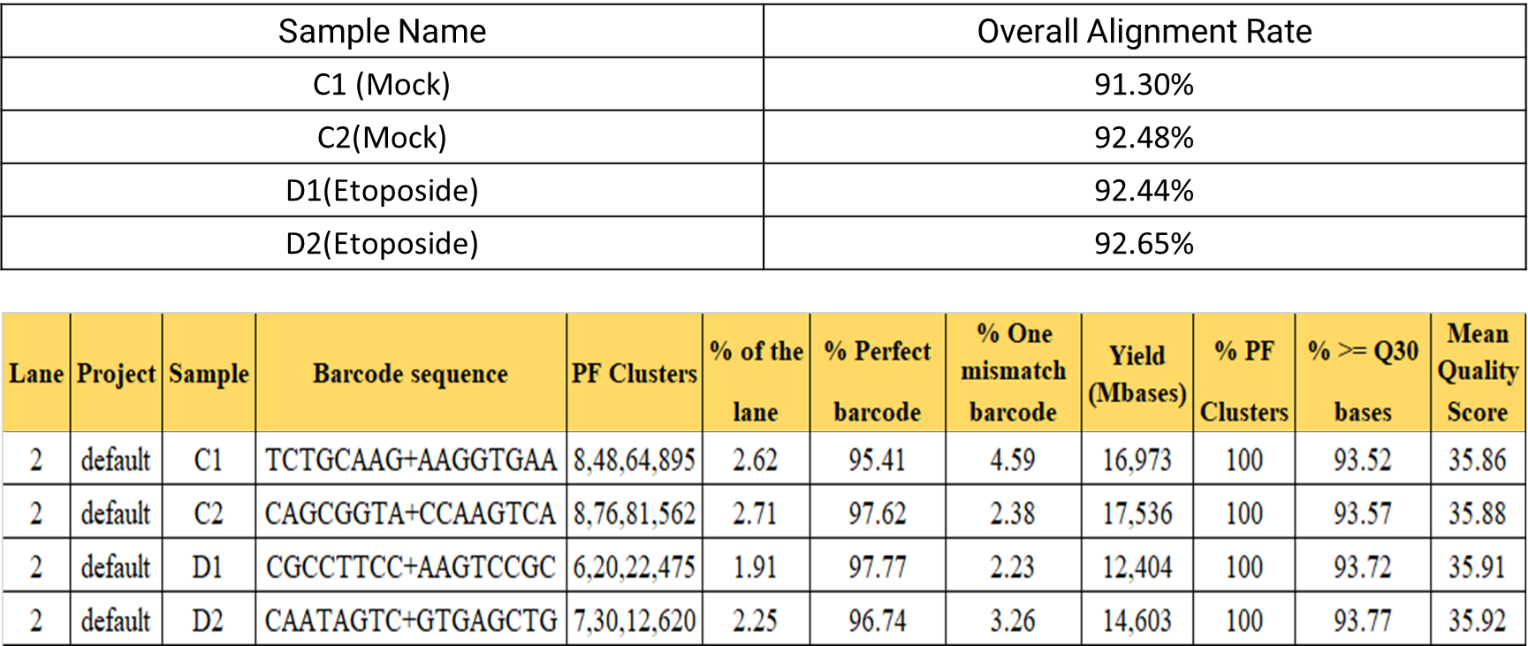

**Figure 3:** Different quality control values and alignment rates of the samples used for RNA Sequencing.

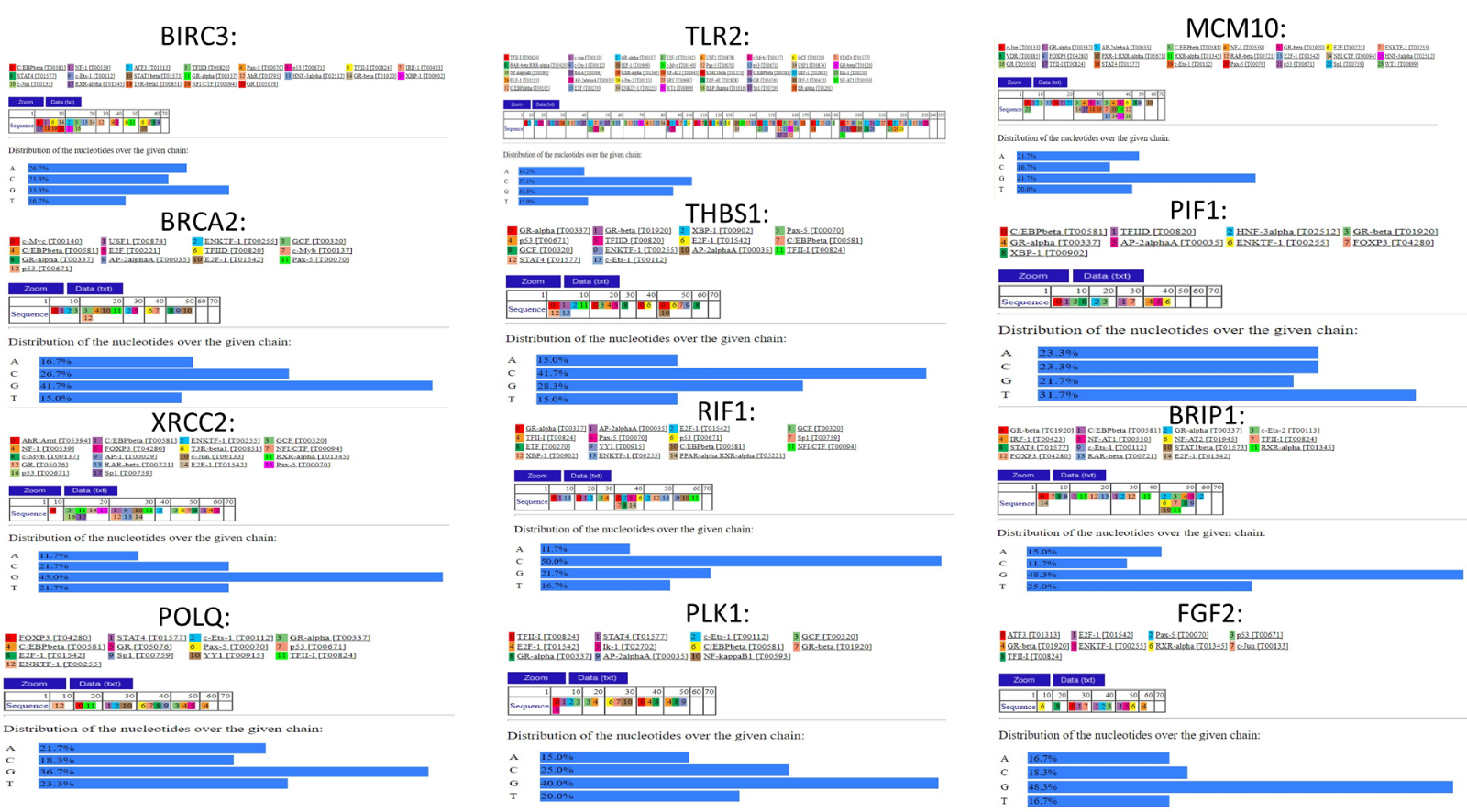

**Figure 4:** The putative transcription factors for each of the 12 gene promoters obtained from PROMO (TRANSFAC ver 8.3)
